# Brain fingerprint is based on the aperiodic, scale-free, neuronal activity

**DOI:** 10.1101/2022.07.23.501228

**Authors:** P Sorrentino, E Troisi-Lopez, A Romano, G Sorrentino, V Jirsa

## Abstract

The possibility to identify subjects from their brain activity was met enthusiastically, as it bears the possibility to individualize brain analyses. However, the nature of the processes generating subject-specific features remains unknown, as the literature does not point to specific mechanisms. In particular, most of the current literature uses techniques that are based on the assumption of stationarity (e.g. Pearson’s correlation), which do not hypothesize any mechanisms, and crashes against a large body of literature showing the complex, highly non-linear nature of brain activity. In this paper, we hypothesize that intermittent moments when large, non-linear perturbations spread across the brain (defined as neuronal avalanches in the context of critical dynamics) are the ones that carry subject-specific information, and that contribute the most to identifiability. To test this hypothesis, we apply the recently-developed avalanche transition matrix (ATM) to source reconstructed magnetoencephalographic data, as to characterize subject-speficic fast dynamics. Then, we perform identifiability analysis based on the ATMs, and compared the performance to more classical ways of estimating large-scale connections (which assume stationareity). We demonstrate that selecting the moments and places where neuronal avalanches spread improves identifiability (p<0.0001, permutation testing), despite the fact that most ot the data (i.e. the linear part) are discarded. Our results show that the non-linear part of the brain signals carries most of the subject-specific information, shading light on the nature of the processes that underlie subject-identifiability. Borrowing from statistical mechanics, a solid branch of physics, we provide a principled way to link emergent large-scale personalized activations to non-observable, microscopic processes.

## Introduction

Higher cognitive functions rely on the complex, coordinated interactions among multiple brain areas. Whole-brain functional measurements (e.g., fMRI, M/EEG) have been used to characterize such interactions, typically using the pair-wise statistical dependencies between regional signals as a proxy (*1*). The time-averaged patterns of interactions (estimated via the Pearson’s or Spearman’s correlation coefficient (PCC, SCC, respectively) in fMRI or via synchronization metrics (among others) in M/EEG) form the static functional connectome (sFC) (*2*). sFCs contain subject-specific information, constituting a “brain fingerprint” which allows subject-identification (*3, 4*).

However, large-scale brain activity reconfigures itself over time (*5*). A recent fMRI-based study used a sliding-window approach to show that specific time intervals carry the vast majority of the subject-specific information (*6*). This finding poses a challenge on the nature of the information upon which subject identification is based. In fact, the intermittent nature of the moments of “high-identifiability” might be generated by non-stationary processes (*7*).

Converging evidence shows that large-scale brain activity has an aperiodic, scale-free component, which mainly drives static connectivity (*8*). The presence of scale-free bursts is not expected for stationary processes, whereas it can be a manifestation of a dynamical system operating in a near-critical regime (characterized by a branching ratio ~= 1) (*9*). Borrowing from statistical mechanics, these scale-free bursts have been defined as “neuronal avalanches” (*10*). Neuronal avalanches are fine-tuned, present across species, across temporal and spatial scales, and have been observed with multiple techniques (*11*). In the human brain, neuronal avalanches spread across gray matter regions through the white-matter bundles linking them (*12*). Furthermore, altered avalanche dynamics is related to clinical disability in multiple neurological diseases (*13*), demonstrating the clinical relevance of this phenomenon.

Here, we hypothesize that neuronal avalanches are the expression of subject-specific, large-scale brain processes. As a consequence, we expect subject-identification based on avalanches to perform better as compared to using the entire available data (for instance, using the SCC). To test our hypothesis, we used data from a previously published cohort (*14*), consisting of source-reconstructed MEG data from 44 healthy subjects. Each participant underwent two magnetoencephalography (MEG) scans, separated by a ~ 1.5 minute-long pause. To identify avalanches, each source-reconstructed signal was z-scored (across-time) and then set to 1 if above a threshold, and to 0 otherwise. Each avalanche was defined as starting when at least one region was above threshold, and as finishing when no region showed unusually high activation. Note that the higher the threshold, the least amount of data is taken into account for identification. For each avalanche, the *ij*^*th*^ entry of the transition matrix contains the probability of region *j* being recruited at time *t*+1, given that region *i* had been recruited at time *t*. We then averaged, edgewise, all the avalanche-specific transition matrices and, after symmetrizing, we obtained per each session a subject-specific avalanche transition matrix (ATM), which captures the spatio-temporal dynamics of neuronal avalanches in a given subject. Then, we used the correlations between the ATMs as a metric of similarity (between the first and the second MEG scans), and used it to define individual “identifiabilities” (i.e., if two ATMs from one subject resemble eachother more than they resemble the ATMs from the other participant, the subject is correctly identified). The average of the individual identifiabilities was defined as the success rate (SR).

First, we set out to define the optimal threshold to maximize identifiability. Under the hypothesis that the spreading of aperiodic, scale-free perturbations carries subject-identifying information, we expect maximal identification to occur when focusing on the scale-free activity, as opposed to taking the whole data into account. Hence, we varied the threshold used to define a region as active. We hypothesize the SR to peak when linear activity is discarded and scale-free activity is preserved.

Then, we compared the identifiability obtained with our approach with three alternative approaches (Fig.1, A). First, we performed identifiability analysis based on connectivity defined as the pariwise Spearman’s correlation coefficients (SC) on the whole dataset. Given the hypothesis that information-carrying interactions occur on the large-scale during neural avalanches, we reasoned that identifiability should be worse using SC as compared to ATM, despite the fact that SC exploits the whole data while the ATMs are only based on a few locations and time points (i.e., the specific moments when an avalanche has recruited specific regions). Then, we selected the moments when neuronal avalanches were occurring, and concatenated the corresponding z-scored time series (i.e., before binarization). Based on this (small) part of the original data, we performed identifiability analysis. In this case, we expect worse identifiability as compared to the ATM, but comparable to the SC based on the whole signal. Finally, we have repeated the same procedure, this time by randomly selecting moments of the z-scored time series when no-avalanches were occurring, and concatenating them to obtain, at each interaction, the same duration as the total duration of all the avalanches. We expect the lowest identifiability in this case (in fact, according to our hypothesis, we would have excluded the most informative parts of the signal from our analysis). We repeated this procedure 1000 times, hence providing a null-distribution representing the SRs to be expected when selecting random segments of the data, under the null hypothesis that avalanches do not carry most of the subject-specific information. An overview of the procedure is provided in Fig.1, B.

**Figure 1.**
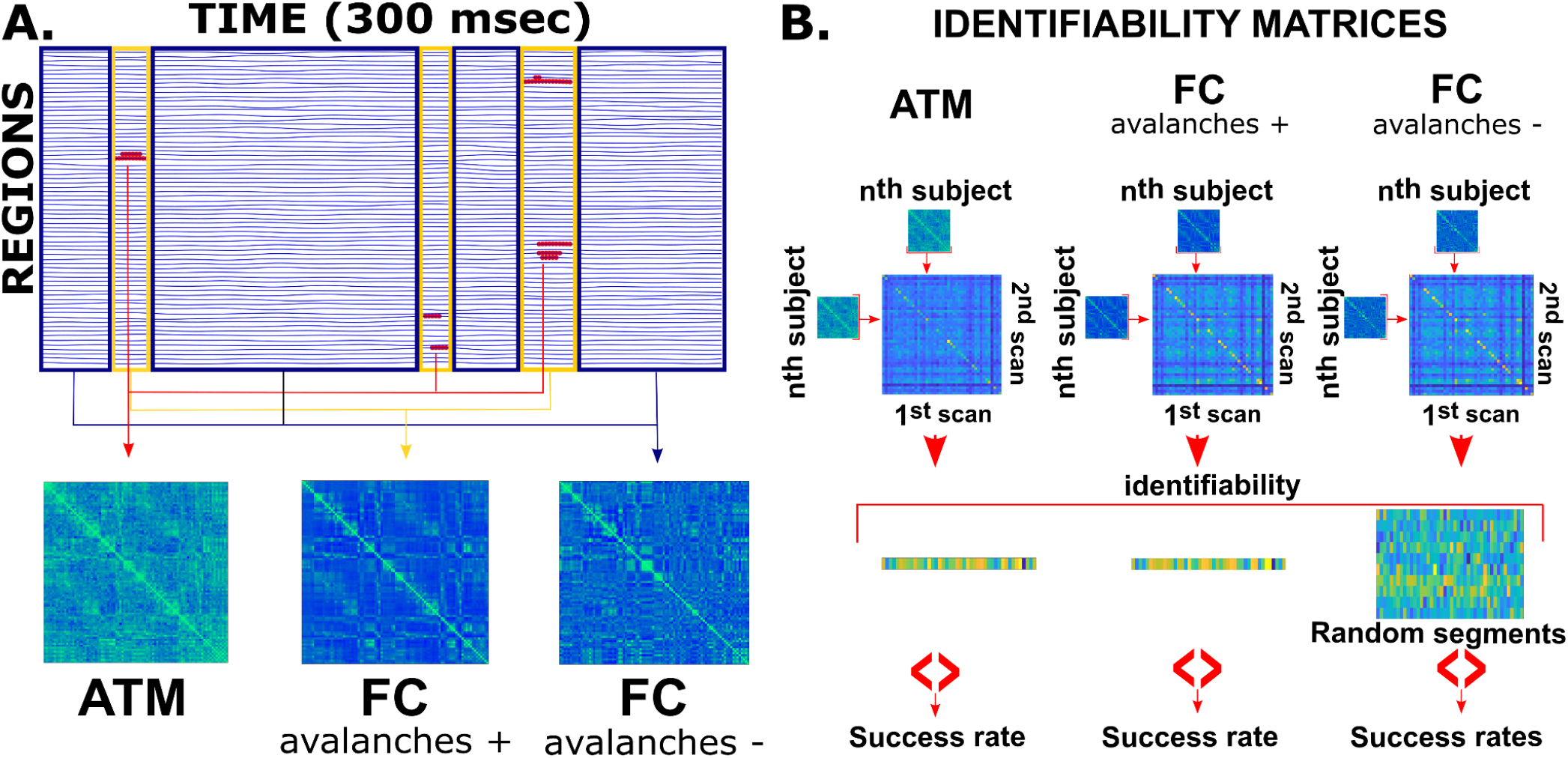
**A**. Data selection. Blue lines represent z-ecored source-reconstructed time-series. In red, the data points that are above threshold. The transitions between these points are the base to compute the ATM. The segments of the data when avalanches are occurring are surrounded by yellow rectangles. All these segments are concatenated, and the Spearman’s correlation is computed in the “FC when avalanche” case. Segments without avalanches are surrounded by blue rectangles. These constitute the vast majority of the data. These segments are concatenated multiple times, each time with a random selection of them, until the same length of the “avalanche” segment is reached. **B**. Construction of the identifiability matrices, containing the Spearman’s correlation coefficient between the connectivity matrix of the first and the second acquisitions. For subject *i*, the success rate is defined as the number of participants that are more similar to subject *i* as compared to subject *i* itself. The success rate refers to the average of the subject success rates.

## Results

In this paper, we set out to quantify subject identifiability analysis using the success rate (SR), defined as the average individual success rates (which captures correctly identified subjects). We have explored the SR as a function of the threshold used to binarize the time-series. This was done under the hypothesis that avalanches define the most “informative” moments with respect to subject-specific activities. If this were not the case, identifiability should drop as the threshold grows, provided that a higher threshold discards more data (note that a z-score of 2.6 discards ~ 99% of a z-score distribution). Our results show that the maximal SR is obtained with a threshold equal to z>|2.6| (Fig 2, A, green line). The identification rate improves as the threshold grows from |1.5| to |2.6|. Hence, the identification rate improves despite the fact that less data is being taken into account, in accordance with the hypothesis that subject identification is based on the spatio-temporal dynamics of neuronal avalanches (which would be, in turn, the main driver of functional connectivity).

**Figure 2.**
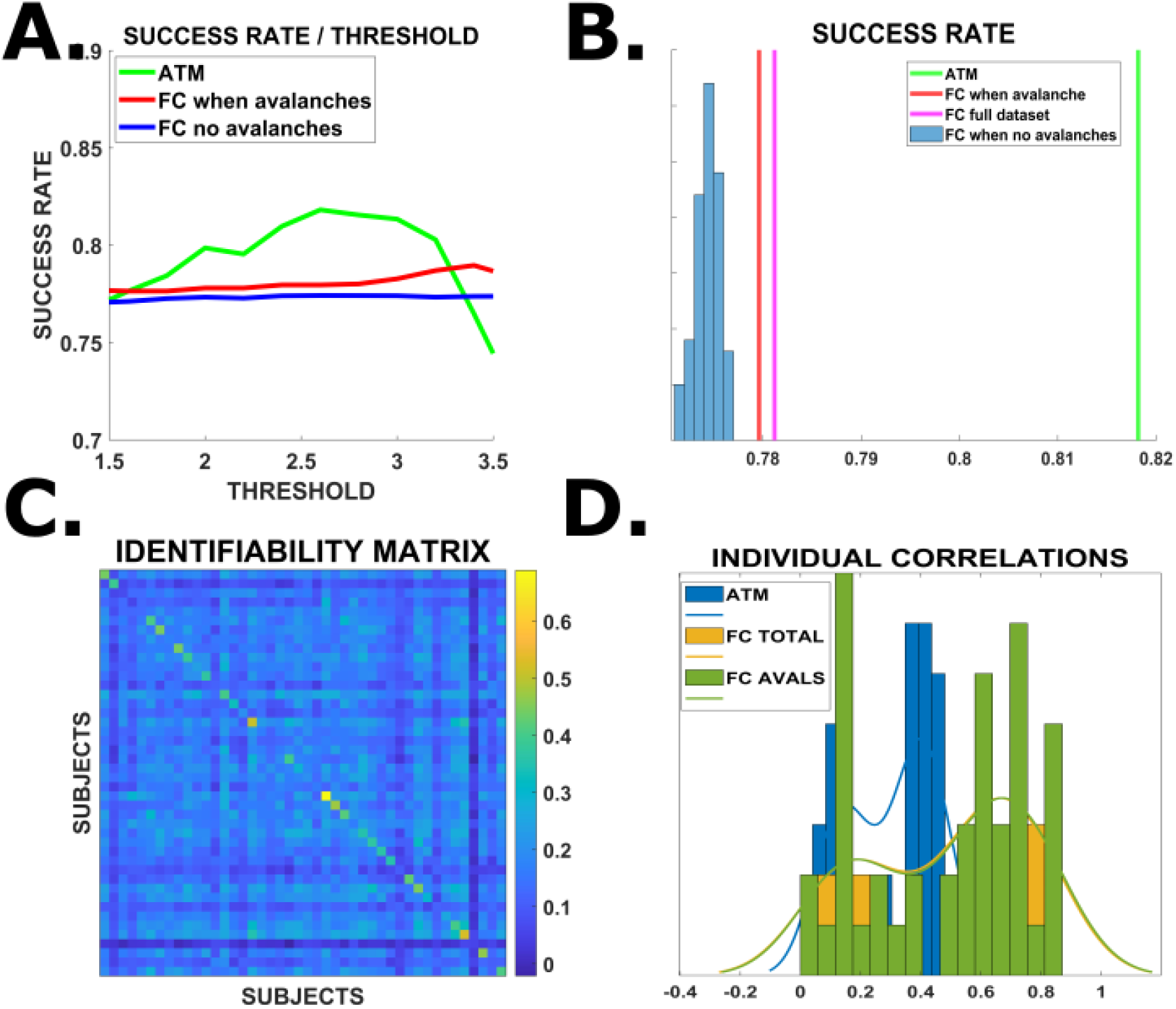
**A**. Success rate as a function of the threshold defining avalanches.The green, red and blue lines refer to the ATM, the FC when there are avalanches and the FC in moments when no avalanches are present. **B**. for the threshold set to z=|2.6| (i.e. the threshold corresponding to the best identification rate), the green, magenta and red lines correspond to the SR obtained using ATM, FC computed on the whole dataset, and FC computed only in moments when there were avalanches, respectively. The distribution in blue refers to 1000 SRs obtained for 1000 random selections of moments when no avalanche was present, each one of the same length of the moments with avalanches. **C**. Identifiability matrix, containing the Spearman’s correlation coefficient between the matrices belonging to first and second acquisitions. The main diagonal contains the similarities of each subject with itself. **D**. The histograms and corresponding distributions refer to the values of the correlation between the first and second acquisitions for each individual. The blue, green and yellow distributions refer to the similarities between the ATMs, the FC based on moments when avalanches are present, and the FC when no avalanche is present, respectively.

Then, for each threshold, we selected from the original z-scored time series the timepoints when avalanches had been occurring (see Fig.1, panel A). Based on these time points, we have computed the adjacency matrices using the SC and used these, instead of the ATMs, to perform subject identification. In other words, we are now using the moments when avalanches were occurring, but we are not selecting the specific regions that were recruited by each avalanche (as we consider all the regions instead). Using this approach, we show that the SR is lower using the SC as compared to using the ATMs (Fig.2, panel A, the red line indicates, for each threshold, the SR based on the SC, while the green line denotes the SR obtained using the ATMs). This might be compatible with the idea that large-scale interactions occur at intermittent time-points and in specific brain regions.

Finally, for each threshold, we have randomly selected the same number of timepoints, but this time choosing among the moments when no avalanche was occurring. Based on this choice, we have again performed subject-identification. We show that, despite the fact that we consider the same amount of data, the identification is worse if neuronal avalanches are excluded from the analysis (Fig.2, A, blue line). In fig.2, B, we report the SR based on ATM (green), on SC based on avalanches (red), on SC based on the whole data (pink), and on the 1000 instances of the SC computed in moments when no avalanches were occurring (blue distribution). This shows that, using SC, the best estimates are obtained from the moments when avalanches are occurring, and this is unlikely to be obtained given a random selection (p<0.001). In Fig.1, C, we reported the similarities between ATMs based on the first (rows) and second (columns) acquisitions. Higher values on the main diagonal denote successful identification.

Then, we moved on to compare the similarities (correlation) between the adjacency matrices based on the SCs (when avalanches were occurring) and those based on the ATMs. Fig.2, D shows the distribution of the correlations for the ATMs (blue), for the SC computed on the whole data (orange), and for the SC computed on the moments when avalanches were occurring (green). It is easy to see that the ATMs show lower correlation values, as compared to SC. However, as shown, higher correlations do not translate into better identifiability, showing that higher correlations might not be driven by subject-specific information.

## Discussion

In this manuscript, we test the role of aperiodic, scale-free, higher-order activations (i.e. Neuronal Avalanches) in subject identifiability. Avalanches are defined as occurring when regions deviate from their baseline activity, which is defined, in turn, on the standardized score of the regional signal. In other words, we select the moments and locations (regions) which show unexpected levels of activity (given the baseline, linear activity of each region), thereby focusing on higher-order activity.

We use the recently described ATM (“avalanche transition matrix”) to capture the spatio-temporal structure of avalanches (*14*). The avalanche-specific transition matrix contains, in the *ij*^*th*^ position, the probability that region *j* will show unusual high activity after region *i* did. In other words, the transition matrix contains the probability that a “wave” of higher-order activations would propagate from *i* to *j*. Averaging across avalanche-specific transition matrices, we obtained one subject- and session-specific avalanche transition matrix (ATM), and utilized the similarity between session-specific ATMs to perform subject identification. We show that fingerprint analysis performed on ATMs improves performance as compared to using matrices based on the Spearman’s correlation coefficient, despite the fact that significantly more data is considered in the latter case. We have replicated the results using Perason’s correlation instead of Spearman’s, and the results are confirmed (not shown). In fact, by construction, avalanches only select the rare, fat-tailed part of the regional activity, and entirely discard the vast majority of data points. However, these few, scattered moments and locations are the ones carrying most subject-specific information. This might talk to the converging evidence showing that aperiodic, scale-free dynamics conveys physiologically meaningful and subject-specific activity (*8, 10, 11, 13, 15*). Accordingly, the spreading of neuronal avalanches, at the individual level, is related to the individual structure of white-matter bundles (*14*). This suggests the idea that an input received by a region (from the rest of the brain) might affect the local processess, provoking an “unusual” activation (*16*). While purely speculative, one could frame this within the “communication through coherence” hypothesis, whereby the incoming inputs would “entrain” local neuronal populations, momentarily favoring their synchronization, and resulting in a signal amplitude that would have not been generated in the absence of the incoming signal (*17*). As said, once such perturbation is generated, it spreads preferentially across the white-matter bundles. Recently, a mechanistic, mean-field based model of the brain dynamics showed that avalanche-like activity might be generated when regions are realistically coupled (*18*). In conclusion, our data show that ATMs provide a mathematically and physiologically rooted, yet straight-forward, way to describe subject-specific fast dynamics.

The correlation values between the SC-based adjacency matrix from the two sessions were generally significantly higher than those obtained using the ATMs. This is somewhat unsurprising, considering that the former exploits the whole dataset, while the latter only a few data points. However, the SC is outperformed by the ATM in terms of subject identification. Hence, one might interpret that as the SC capturing features that are shared by the whole dataset, while the ATMs focusing more specifically on subject-specific dynamics. Furthermore, the ATMs do not consider zero-lag associations, hence correcting for field spread, while SC is biased in that sense (*14*). Connectivity metrics that do not correct for volume conduction tend to outperform, in terms of fingerprinting, those who do. This is interpreted as an effect of the contribution of volume conduction to identifiability, provided that the geometry of the head shapes the field-spread, and that this carries subject-specific information, although not neurophysiological in nature (*19*). In this sense, the higher similarities obtained by the SC might not be due to similar patterns of activity but, rather, by similar structural features. In conclusion, ATMs are effective in selecting the locations and moments where large-scale patterns spread, which might be directly related to subject-specific neurophysiological mechanisms. In fact, when SC is performed on the data segments that contain the neuronal avalanches, the performance remains fairly high, confirming that the avalanches indeed represent the moments where subject-specific dynamics is manifesting itself. Accordingly, when subject identification is performed based on moments when the avalanches are not occurring, the most informative part of the signal is lost, and the performance never reaches the levels observed when avalanches are included.

In conclusion, ATMs appear as a straight-forward, yet principled way to select the data that convey subject-specific, large-scale spatio-temporal dynamics. This opens new venues to characterize subject-specific brain dynamics in health and disease, and provides new observables to tune subject-specific brain models. Furthermore, it corroborates the idea that large-scale interactions are of higher-order (i.e. non expected by a linear process), and should be treated as such when processing brain signals. The ATM is a useful tool to quantify such higher-order large-scale dynamics. Finally, task-based studies might be analyzed using ATMs, in order to identify the role of specific regions and/or edges in specific behavioral functions.

## Methods

### Cohort, data acquisition, preprocessing, source-reconstruction, avalanches estimation

Methods for the description of the cohort and the data acquisition, preprocessing, source-reconstruction, avalanches estimation and avalanche transition matrices have been described in detail in (*14*). In short, we recruited 58 young adults (male 32 / female 26, mean age ± SD was 30.72 ± 11.58), right-handed and native Italian speakers with no major internal, neurological or psychiatric illnesses and no use of drugs or medication that could interfere with MEG/MRI signals. The study complied with the Declaration of Helsinki and was approved by the local Ethics Committee. All participants gave written informed consent. 3D T1-weighted brain volumes were acquired at 1.5 Tesla (Signa, GE Healthcare) after the MEG recording (*14*). The MEG registration was divided in two eyes-closed segments of 3:30 minutes each, separated by ~1 minute long break. To identify the position of the head, four anatomical points and four position coils were digitized. Electrocardiogram (ECG) and electro-oculogram (EOG) signals were also recorded. After an anti-aliasing filter, MEG signals were acquired at 1024 Hz, then a fourth order Butterworth IIR band-pass filter in the 0.5-48 Hz was applied. Noisy channels were identified and removed manually, Principal Component Analysis and supervised Independent component analysis were used to remove environmental noise and physiological artifacts, respectively. 47 subjects were selected for further analysis. The time series of neuronal activity were reconstructed in 116 regions of interests (ROIs), based on the Automated Anatomical Labeling (AAL) atlas, using the Linearly Constrained Minimum Variance (LCMV) beamformer algorithm, and the native structural MRIs. The sources were reconstructed for the centroids of each ROI (*20*). Finally, we considered a total of 90 ROIs for the AAL atlas, since we excluded the cerebellum because of its lower reliability in MEG (*6*). All the preprocessing steps and the source reconstruction were made using the Fieldtrip toolbox (*21*). To study the dynamics of brain activity, we estimated “neuronal avalanches” from the source-reconstructed time series. Firstly, the time series of each ROI was discretized calculating the z-score, and then identifying positive and negative excursions beyond a threshold. The value of the threshold was varied between 1.5 and 3.5, to spot the value that maximized identifiability. A neuronal avalanche begins when, in a sequence of contiguous time bins, at least one ROI is active (i.e., above threshold), and ends when all ROIs are inactive (*13*). The branching parameter was geometrically averaged across avalanches and subjects. In fact, systems operating at criticality typically display a branching ratio ~1. In detail, the branching ratio was calculated as the geometrically averaged (over all the time bins) ratio of the number of events (activations) between the subsequent time bin (descendants) and that in the current time bin (ancestors) and then averaging it over all the avalanches, as:

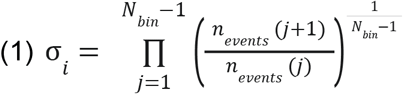

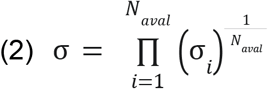

Where σ_*i*_ is the branching parameter of the i-th avalanche in the subject, Nbin is the total amount of bins in the i-th avalanche, Naval is the total number of avalanches in the dataset. The results shown are derived when taking into account all avalanches. However, we repeated the analysis taking into account only avalanches longer than 5 time bins, as well as only avalanches longer than 20 time bins (this furhter reducing the selected datapoints), and the results were unchanged (see supplementary material 1). Then, an avalanche-specific transition matrix (TM) was calculated, where element (*i, j*) represented the probability that region *j* was active at time *t+*, given that region *i* was active at time *t*, where ~3ms. The TMs were averaged per participant (i.e. across avalanches), and then per group, and finally symmetrized. The introduction of a time-lag makes it unlikely that our results can be explained trivially by volume conduction (i.e. the fact that multiple sources are detected simultaneously by multiple sensors, generating spurious zero-lags correlations in the recorded signals). Please also refer to (*14*), response to reviewers, for extensive analysis on field spread on ATMs in this dataset.

### Fingerprint analysis

Fingerprinting analysis was performed similarly to (*22*). In short, for each subject, for each of the two acquisitions, an adjacency matrix was built (either the ATM or the ones based on the SC). Hence, each participant had two symmetric matrices, each referring to one of the two acquisitions. Pearson’s correlation coefficient between the vectorized lower triangular matrices of the two acquisitions were used as a metric of similarity. Hence, an identifiability matrix was built (Figure 2, panel B), with rows and columns referring to the first and the second acquisition for each participant, respectively. Hence, the main diagonal contains the similarity between each subject with themselves in the first and the second acquisition. The off-diagonal elements contain the similarities of each subject with every other subject. Hence, if, for one subject, the value in the main diagonal is the highest, that subject is correctly identified. The individual success rate was defined as the number of participants that were more similar to a subject as compared to how similar the subject was across the two sessions. The average of the individual success rates was defined as the success rate, which refers to the whole population, and was used as a readout in our analysis. Our analysis was repeated using Pearson’s correlation instead of Spearman’s (despite the fact that the signal is non-linear), and the resutls are confirmed.

## Supplementary materials 1

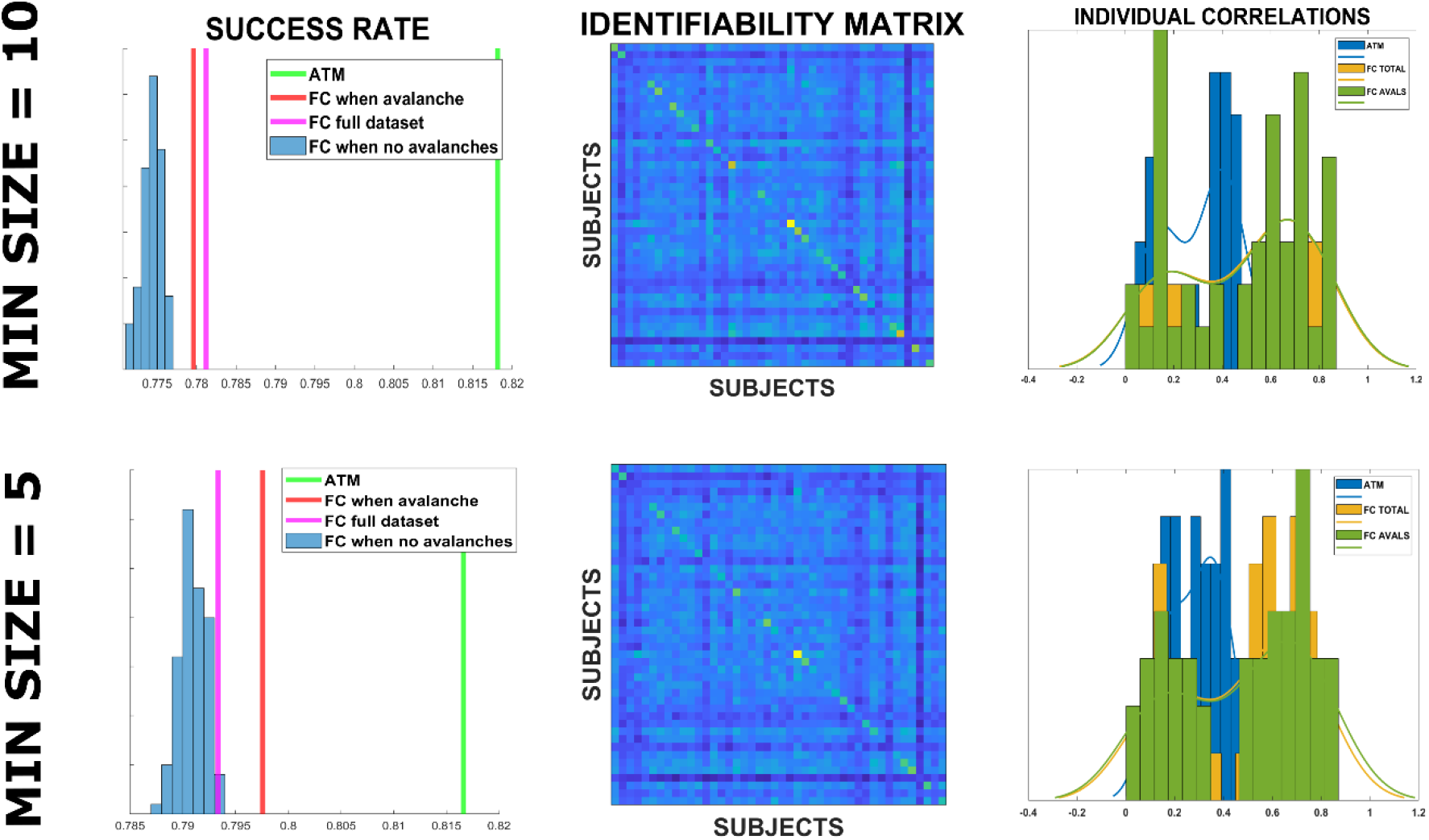

The figure shows the resutls replicated changing the lower bound for the size of avalanches to be included in the analyses, i.e. 10 time bins (top) or 5 time bins (bottom).

